# Chasing amylase: genotypic diversity in *Geukensia demissa* may be environmentally driven

**DOI:** 10.1101/2025.07.21.665945

**Authors:** P.N. White, C. Ewers, J.P. Wares

## Abstract

Prior results suggested an unusual sequence polymorphism in an amylase gene region of the ribbed mussel *Geukensia demissa*. We used targeted sequencing, spatial analysis of polymorphism data and a high-quality *de novo* genome assembly to assess the likelihood that this diversity is inherited as a single Mendelian locus. Our results suggest this is likely, and that the frequencies of the dominant allele groups change dramatically along the Atlantic coast of the United States, particularly across Cape Hatteras. The single-copy nature of the sequence being studied was verified through the development of a new reference genome for *Geukensia demissa*, described here. Although the overall pattern of sequence diversity is atypical, with much higher allelic divergence than is expected given overall genomic diversity, we believe this may be an ecologically valuable molecular marker for study in ribbed mussels.

## Introduction

The ‘molecular’ study of biodiversity has gone through a number of significant changes; throughout, researchers have tried to understand the mechanisms - both neutral and adaptive - that lead to spatial or temporal patterns in that biodiversity. These mechanisms can be very difficult to disentangle (Meirmans 2012, Wang and Bradburd 2014). Often, our focus is on how the environment plays a role in adaptation, and a mistake is made by overlooking purely neutral mechanisms that drive at least part of the pattern. Even when there are clear signatures of adaptation to distinct environments, our ability to resolve the genomic contribution to that adaptation can be limited by available data (Moyle et al 2012). Paradoxically, the more genomic data we are able to collect, the likely proportion of those data that are truly adaptive shrinks (Rellstab et al. 2015) because most of the genome is merely responding to demographic forces (Wares et al. 2021, Duffin et al. 2026).

With that in mind, early studies had few genomic markers to work with, often using allozyme markers, and these strong metabolic drivers of environmental tolerance were likely to be highlighted in such studies (Marden 2013). This is certainly true for studies of the evolutionary ecology of marine invertebrates; Hilbish & Koehn (1985) summarized key work on variation in salinity tolerance in the mussel *Mytilus edulis* showing that products of the leucine-amino-peptidases (*Lap*) gene are mechanistically involved, and that natural selection drives the cline in allele frequencies at this locus for this species. Similarly, Schmidt and Rand (2001) identify microhabitats separated by only fractions of a meter in the intertidal – but with very different exposure to heat and desiccation – that result in distinct genotype frequencies at the mannose phosphate isomerase (*Mpi*) locus in the barnacle *Semibalanus balanoides*.

This classic work in *S. balanoides* has recently been supported by genomic evidence (Nunez et al 2021) showing that balancing selection across the spatial scale of many species may be key in population persistence across such large ranges of physiological requirements. Still other genome-scale studies can now clarify which discrete parts of the genome drive environmentally-relevant patterns of diversity most effectively (Bay et al. 2018, Eppley et al. 2026). With each advance in available genomic resources for a species, there are new opportunities to find those few regions that lead to insight about adaptation.

With such challenges in mind, Erlenbach & Wares (2023) evaluated patterns of diversity and divergence from a subset of commonly-studied metabolic loci in the ribbed mussel *Geukensia demissa*, a mussel common in East coast saltmarshes of North America from the Canadian maritimes to the Gulf of Mexico. The sequence diversity for each individual was reconstructed from RNA sequence data, using a *de novo* transcriptome assembly as the scaffold for the sequences recovered in individuals from two distinct spatial populations (Georgia and Massachusetts, U.S.A.). While most of the loci were effectively panmictic between these “northern” and “southern” range mussels, the most divergent metabolic transcript identified between these populations was for an amylase gene region, with statistically highly significant sequence divergence (Hudson’s Snn 0.947*) and a positive value of Tajima’s D across all samples, suggesting a non-random pattern of evolutionarily divergent sequences (Ewers and Wares 2012). This gene region appeared to be more divergent between sites than even mitochondrial sequences (Díaz-Ferguson et al 2009, Erlenbach & Wares 2023), suggesting a spatial pattern that could be of value in studying ecotypic diversity in *G. demissa* across several environmental gradients on the Atlantic coast of North America.

If the samples in Erlenbach & Wares (2023) had incidentally recovered distinct copies of an amylase gene family, our comparison across spatial scales would not be one of homologous evolutionary divergence but of a misunderstanding of the genome of ribbed mussels. Yet, if the diversity behaves as a single locus, then the spatial patterns in genotype frequency might highlight correlated metabolic variation along the environmental gradients that *G. demissa* spans. Recognizing that even with a well-scaffolded genome there are a number of ways in which the bioinformatic inference of Erlenbach & Wares (2023) could be misled through missing data or indirect annotation (Whibley et al. 2021), we have resampled *G. demissa* along the Atlantic coast to generate sequence data at the amylase locus to study the spatial patterns and likely inheritance mode of the observed diversity. We then also compare our results with a newly-assembled genome for *G. demissa* as well as other confamilial genomes to solidify expectations for exploration of this marker.

Amylase itself is of great interest for studying the ecophysiology of a filter feeder like *G. demissa*, as the primary diet of these mussels is starch-rich phytoplankton. While phytoplankton availability varies latitudinally along this coast along with sea surface temperature, amylase diversity has also been shown to have significant role in growth in oysters (Prudence et al 2006) and food choice in marine amphipods (Guarna & Borowsky 1993, Rodriguez-Viera et al., 2021). The sequence diversity of the primary ‘types’ of amylase recovered in Erlenbach & Wares (2023) suggested significant divergence between the two sequence classes, including nonsynonymous amino acid substitutions. It is thus plausible that the distinct forms have distinct metabolic properties. If the diversity we can examine for amylase in *G. demissa* behaves as a single locus and varies corresponding to environmental gradients, it merits further attention as possibly ecologically important gene region (Skovmand et al 2018). With these goals in mind, we present sequence and genotype diversity for amylase in ribbed mussels, along with a new reference genome, to evaluate the utility of this genomic marker for further ecotypic and biogeographic analysis in this species.

## Methods

We assume that for a multi-copy gene family, either 2 or more independent copies will consistently be recovered using PCR (in which case we would infer an excess of heterozygous individuals relative to Hardy-Weinberg expectations), or that the primers only amplify a single locus within that gene family because of differences in primer region diversity. To assess whether our focal amylase in *G. demissa* is a single copy gene region, we assay the recovered diversity across multiple individuals from multiple locations. Based on the sequence data reconstructed in Erlenbach & Wares (2023), we aligned the amylase sequence region from Massachusetts and Georgia individuals and developed 2 sets of PCR primers. While one primer pair generated inconsistent PCR results – later examination showed that the targeted region spans an intron-exon boundary – the second pair GdAmy239F (CCATCTTATTGACATTGGTGTAG) and GdAmy593R (TCCTCTCTGGTTGTCATGGTT) amplify a 354bp fragment when using an annealing temperature of 50°C (95° 3:00, then 35x (95° 0:30, 50° 0:30, 72° 1:00), 72° 5:00).

Table 1 indicates which samples were subsequently sequenced. Amplicons were cleaned using an exonuclease protocol as in Wares (2024) and submitted for bidirectional sequencing (psomagen.com). End trimming of sequence data used default settings in CodonCode Aligner. All nucleotide calls with PHRED scores <15 were visually assessed and called as unknown, an IUPAC ambiguity, or as a single base. Paired sequence data were cleaned and aligned using CodonCode Aligner (v 11.0.1), retaining IUPAC ambiguity codes at the sites of confirmed within-individual polymorphism (roughly equal height of the two chromatograph peaks at a site). Consensus sequences for all individuals were then aligned to each other, confirming the consistent location of polymorphic sites in the data. Individuals were thus called as homozygotes for either the “northern” or “southern” allele recovered in sequences from Massachusetts and Georgia, respectively in Erlenbach & Wares (2023), or heterozygotes when that individual appeared to carry both alleles.

**Table 1.** Sample locations and results of Hardy-Weinberg assessment of observed sequence or amplicon data. Data combined for 2 proximal sites in New Jersey (*).

| Site | Lat/Long | Date Collected | n | Data Collected | 'Southern' Frequency | HWE p-value |
| --- | --- | --- | --- | --- | --- | --- |
| Antigonish, Nova Scotia | 45.668°, - 61.878° | 12/6/2023 | 11 | Seq | 0.273 | 0.932 |
| Plum Island, MA | 42.759°, -70.891° | 3/6/2024 | 14 | Both | 0.214 | 0.988 |
| Woods Hole, MA | 41.520°, -70.668° | 3/11/2024 | 20 | Both | 0.219 | 0.982 |
| Goldstar Battalion Beach, NY | 40.897°, -73.436° | 6/1/2024 | 20 | Digest | 0.150 | 0.905 |
| Green Island, NY | 40.617°, -73.504° | 6/2/2024 | 20 | Digest | 0.375 | 0.968 |
| Boat Basin, NJ* | 39.555°, -74.363° | 12/10/2024 | 14* | Seq | 0.214 | 0.963 |
| Bivalve, NJ* | 39.234°, -75.031° | 12/11/2024 | 14* | Seq | 0.214 | 0.963 |
| George Island Landing, MD | 38.041°, -75.363° | 7/25/2024 | 20 | Digest | 0.550 | 0.982 |
| Gloucester Point, VA | 37.307°, -76.417° | 7/24/2024 | 20 | Digest | 0.200 | 0.969 |
| Roanoke Sound, NC | 35.903°, -75.604 | 4/26/2025 | 10 | Digest | 0 | NA |
| Atlantic Beach, NC | 34.702, -76.751 | 4/22/2025 | 20 | Digest | 0.750 | 0.967 |
| Wilmington, NC | 34.175, -77.932 | 5/19/2025 | 10 | Digest | 0.625 | 0.607 |
| Charleston, SC | 32.762°, -79.967° | 10/1/2024 | 20 | Digest | 0.925 | 0.997 |
| Savannah, GA | 32.038°, -81.048° | 9/1/2024 | 18 | Both | 0.972 | 0.999 |

To assess the likelihood of these data deriving from the same genomic locus, the frequencies of each allele were estimated from source populations in Georgia, New Jersey, Massachusetts, and Nova Scotia (see Table 1). From these frequencies, expected genotype frequencies were calculated and compared with observed “genotypes” using a χ^2^ test.

The observed site differences between the northern and southern sequences identified amino acid substitutions within an open reading frame; one set of nonsynonymous variants fall within the cut site motif (GAT|ATC) of *Eco*RV restriction enzyme. Table 1 lists the additional samples evaluated using these fragment length polymorphisms. Digestion reactions included 1µl of *Eco*RV (New England Biolabs) in a reaction containing 2µl NEB 10X buffer, 14 µl water, and 3µl of PCR amplicon, incubated at 37° for 15 minutes. We evaluated the product of this reaction on a 1.5% agarose gel, running at 100V for 45 minutes. Amplicons that are not cut by this enzyme only contained the ‘northern’ sequence, while completely cut amplicons represented the presence of only the ‘southern’ sequence; the resultant fragments are approximately the same length so appear as a single band. The presence of two distinct bands on the gel – of same length as uncut and same lengths as cut – reflect apparent heterozygotes.

We tested this method of assessing genotype and directly compared with the results from dideoxy sequencing data and found a 100% match from samples at Woods Hole, Massachusetts (n=20), validating this approach for the remaining spatial samples of *Geukensia*. As above, all spatial samples were evaluated for their potential deviation from Hardy-Weinberg expectations.

In addition to determining the pattern of apparent genotypic diversity within and across samples, sequence data were separated into probable haplotypes (alleles) using the PHASE algorithm in DNAsp6 (Rozas et al. 2017). These reflected the prior assessment of two primary sequence types in Erlenbach and Wares (2023) exactly. Population-level sequence diversity was thus assessed by estimating π and Tajima’s D by sample location as well as by sequence type (northern or southern) across all locations. Between sample locations, spatial divergence in sequence frequency was assessed using Hudson’s (2000) S_NN_ metric.

The final data for amylase allele frequency by location were compared to variation in local sea surface temperatures (SST) using data from the World Ocean Atlas (Reagan et al. 2024). We chose mean annual SST for the nearest 1/4° graticule to each site averaged over 1991-2020; the mean winter quarter SST; and the mean summer quarter SST. These values were tested with simple correlation to the ‘southern’ allele frequency at each site.

A novel genome assembly was developed by CD Genomics, using PacBio and Hi-C sequencing complemented with RNA sequence data and annotation. The source individual of *G. demissa* was collected in January 2026 from Fort Johnson, South Carolina. High molecular weight DNA was isolated using a standard protocol at CD Genomics. The PacBio Sequel II HiFi reads were filtered using fastplong (Chen 2023), removing low quality bases and reads shorter than 5kb.

Illumina Hi-C data were filtered and clipped using Fastp (Chen et al 2018). The PacBio reads were used for *de novo* genome assembly with hifiasm; this assembly was then processed using purge_dups to remove redundant contigs. Hi-C assisted genome assembly was obtained using the 3D-DNA pipeline (Dudchenko et al. 2017). The resultant scaffolds were inspected using Juicebox Assembly Tools (Robinson et al 2018).

Repeats were identified with RepeatModeler (Flynn et al 2020); RepeatMasker identified known repetitive elements. Soft-masking of the genome assembly, using Funannotate (Palmer and Stajich 2020), generated the final library. RNA sequence data were assembled with Trinity and aligned to the scaffolds using BLAT. Gene prediction was performed using the Funannotate framework; proteins were identified using the UniProtKB/Swiss-Prot database and data from related bivalves.

The resultant sequence data for the amplicon studied here was compared with an independent *de novo* RNA transcriptome for *G. demissa* as well as the chromosome-scale *de novo* genome assembly of *G. demissa* described above. The *de novo* RNA transcriptome was assembled with the Oyster River Protocol (MacManes 2018) using data from individual MAC5, which was the most complete sequence library from Erlenbach & Wares (2023; NCBI PRJNA729970). Our amplicon sequence data were locally aligned against these resources using minimap2 (Li 2018).

## Results

Sequence data for amylase were uploaded to NCBI Genbank (accession numbers PQ672190-PQ672279 and PV013581-PV013594) from the 84 individuals collected throughout the species range. Table 1 shows the results of assessing sequence and sequence combination (i.e. genotype) frequencies observed across all sites including both sequenced and restriction-genotyped individuals; each location generated a very close match between observed and expected genotype frequencies, with all χ^2^ test results showing p > 0.90 other than the small sample from Wilmington NC (p 0.607). Pooling the observations from Atlantic Beach and Wilmington results in a χ^2^ test result with p of 0.868. Genotype data by individuals are available in White & Wares (2026).

Genome-wide nucleotide diversity π was estimated using Illumina sequence data from Wares (2023), though RAD data will typically present a biased estimate of genomic diversity (Arnold et al. 2013). The all-sites VCF files from those data were analyzed using pixy (Samuk & Korunes 2021) using a window size of 100nt, resulting in an average π = 0.00094 for a subset of mussels from northeastern Florida and the Georgia coast. We note this is not a random sample of the genome as it only includes those regions captured by RAD-seq (Arnold et al 2013). In contrast, estimates of π for the amylase sequences in this study are similar within allele groups (0.00048, 0.00124) but an order of magnitude higher when all sequences are analyzed together (π = 0.01436). Further evidence of admixture of distinctly evolved sequences comes from Tajima’s D, which is negative within allele groups (−1.488, −1.148) but positive when all sequences analyzed together (D = 0.378).

Divergence among spatial samples was statistically significant, with Hudson’s Snn of 0.394 (p<0.001). For the paired regional data of Georgia versus New Jersey, Snn is 0.838 (p<0.001); for New Jersey versus Massachusetts, Snn is 0.599 (ns) and for Massachusetts versus Nova Scotia Snn is 0.661 (ns). There is a strong correlation of amylase genotype frequencies with temperature (Figure 1, Table A1); in particular comparison of mean annual SST (r = 0.724) and mean winter quarter SST (r = 0.756), with the correlation to mean summer quarter SST being lower (r = 0.559).

**Figure 1.**
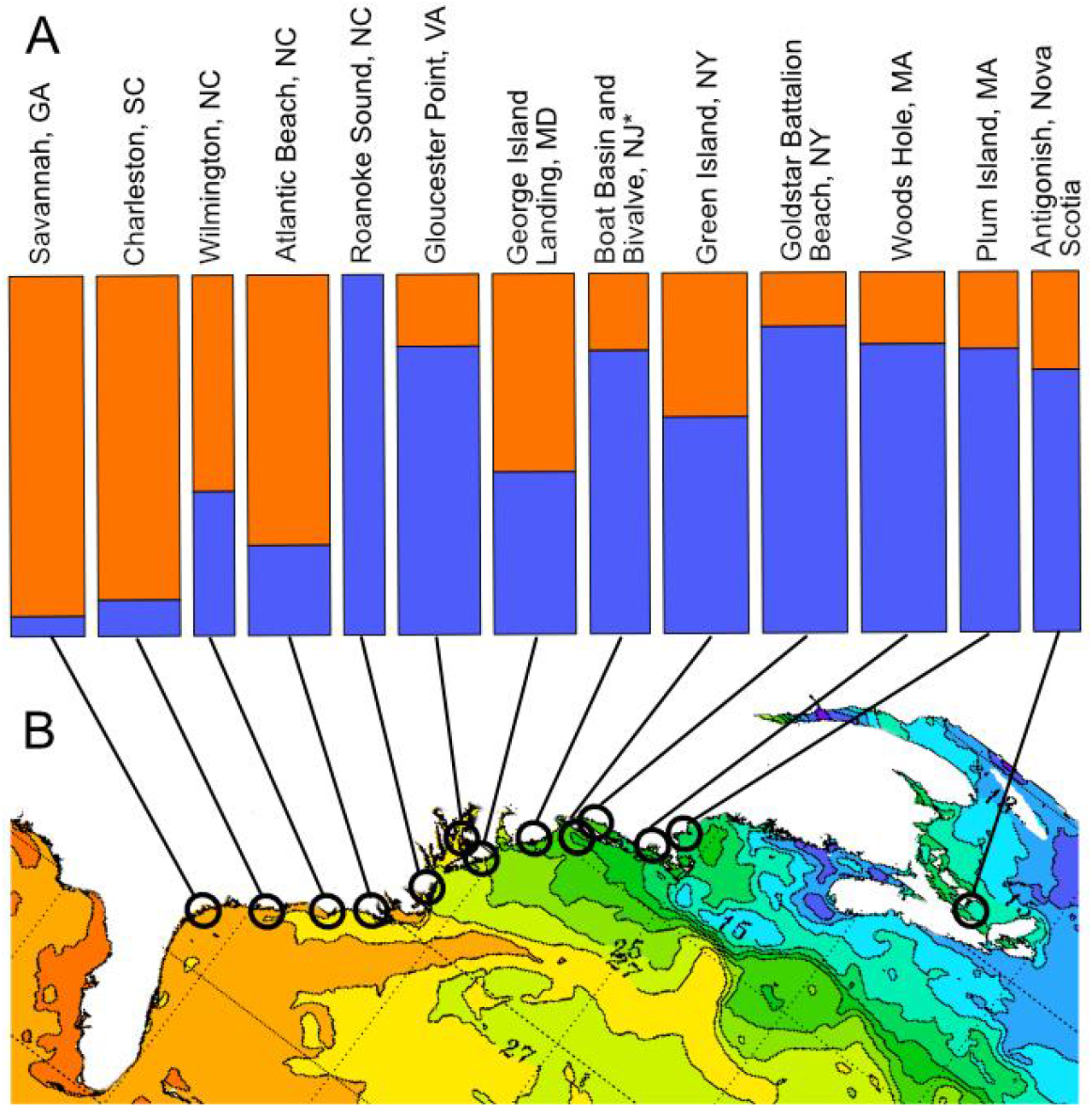
Sampling localities along the east coast of North America (note rotation of map) with (A) relative frequency of southern (orange) and northern (blue) amylase allele and (B) average sea surface temperature (from ospo.noaa.gov/products/ocean/sst). Specific temperature data presented in Appendix.

**Figure 2.**
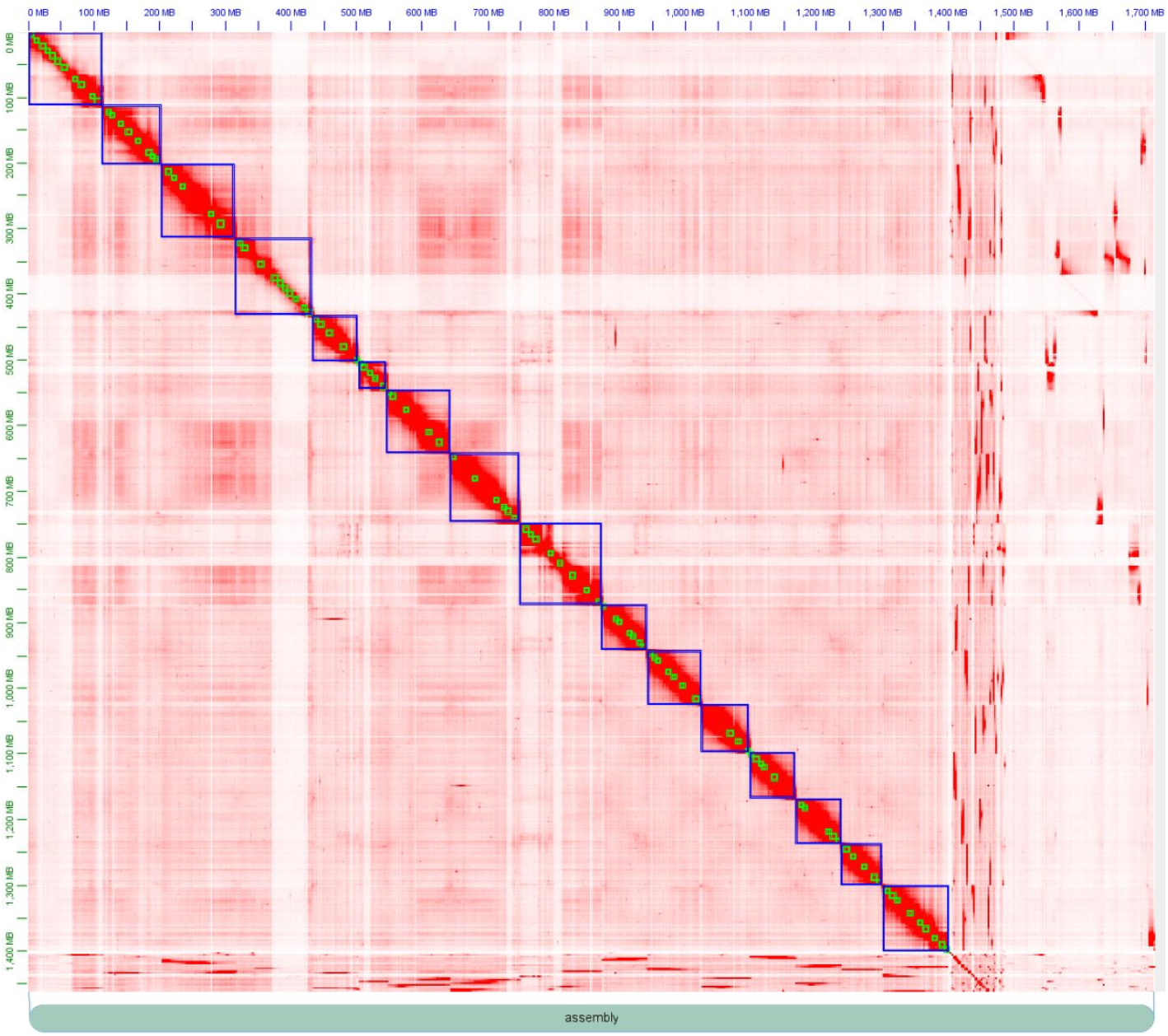
Whole-genome Hi-C interaction map of the assembled genome. Physical interaction frequencies across the genome, as well as scaffold size, support inference of 16 chromosomes. Blue squares delineate the assembled chromosomes, with green squares representing primary contigs. Provided by CD Genomics.

Sequences generated above were compared using BLAST to available genomic resources for *Geukensia* and related bivalves. The *de novo* RNA transcriptome generated from sequence data in Erlenbach & Wares (2023) had a BUSCO score of 93.8% (single copy: 85.5%, duplicate copies: 8.3%) and a Transrate score of 0.374. BLAST against this assembly in Geneious (110,113 sequences) results in the “northern” sequence and “southern” sequence both having a highest pairwise identity with the same single assembled transcript (which itself BLASTs to the nr database at NCBI as “Mytilus californianus alpha-amylase like” XM_052233366, 74.9% identity; Mytilus edulis alpha-amylase like” XM_071286820, 75.1% pairwise identity; or two hits in *Mytilus galloprovincialis* XM_076226570, XM_076226572 with pairwise identities of 75.0% and 74.3%) and no other hits, even when the e-value is increased to 1e-10. The two hits in *M. galloprovincialis* are themselves 90.7% identical.

Completed sequencing of the *G. demissa* genome included 176.6 Gb of PacBIo HiFi reads (~125x coverage), and 243 Gb of HiC sequence data. The data were assembled into a total of 16 chromosomes with another 80 fragment scaffolds and the mitochondrial genome (Figure X); 4 scaffolds were removed through contaminant filtering during processing at NCBI. Scaffold N50 is 96.23Mb, with the largest scaffold at 125.21 Mb. The genome size is 1.41Gb. BUSCO analysis identified 94.3% BUSCO complete genes, 93.5% of those are single-copy. Annotation data are available at X. The final assembly has been deposited in GenBank under BioProject PRJNA1462732.

Alignment of the “southern” and “northern” type amylase gene fragment (acc. nos. PQ672259, g_LTER1000; PQ672226, h_GDWH08) against the complete de novo genome assembly resulted in a single hit on scaffold ptg000142l_1 with pairwise identity of 98-100%, depending on the amylase allele being compared. From additional analysis using minimap2 (Li 2018), this scaffold was generated during the PacBio assembly but purged as a possible duplicate region; the purged scaffolds are also available at BioProject PRJNA1462732. The only mismatches were the known SNPs between “northern” and “southern” type, which were coded with ambiguity scores in the assembly, indicating that the reference genome is heterozygous for amylase. Alignment of the amplicon sequence against our ORP reference transcriptome, which had a BUSCO score of 94.8%, with 88.4% of those single-copy, resulted in a single assembled transcript (TRINITY_DN26139_c0_g2_i1) matching.

## Discussion

The amplicon data generated here for *G. demissa* behave like a single-copy locus in one way (with similar observed/expected genotype frequencies in all samples) yet exhibit extraordinary allelic divergence between ‘alleles’, as might be seen in paralogs or introgressed orthologs (Ewers and Wares 2012, Erlenbach & Wares 2023). The frequency of these alleles and resultant genotypes transitions dramatically along the Atlantic coast distribution. Attempts to BLAST the divergent amylase sequences in a *de novo* RNA transcriptome or the new *G. demissa* genome presented here result in only a single matching contig in each case, and single hits with low identity (because of evolutionary divergence) in confamilial genomes of *Mytilus*.

The higher-quality genome for *Geukensia* is an important resource ensuring the inheritance mode of these sequences, since *Geukensia* and *Mytilus* themselves are more than 300 my divergent (Audino et al 2020).

This new genome for *G. demissa* is of comparable size to those of the genus *Mytilus*, but appears to have 16 chromosomes rather than the 14 identified in *M. edulis* and *M. galloprovincialis*, the species with the highest-quality reference genomes. Our hybrid genome assembly supports the inference from frequency-based comparisons of genotypes across the range of *G. demissa* that the amylase region described here is single-copy. However, this instance of amylase – found from initial long-read sequencing but discarded in the final assembly process – also highlights the value in iterative improvement of reference genomes (Whibley et al. 2021). The value of having a highly divergent genome for the family Mytilidae may be in exploring other changes in genome or chromosome architecture in an ancient and ecologically valuable bivalve family (Corrochano-Fraile et al. 2022).

The overall pattern of amylase diversity in *Geukensia* is now clarified, and at this point appears to be inherited as a single diploid locus. We cannot identify any other mechanism in which the types are co-amplified in frequencies consistent with Hardy-Weinberg expectations within regions, while varying across regions. As noted, the two allele groups are quite divergent with little segregating diversity other than the identifying substitutions between them. Though balancing selection may be more common than once thought in marine intertidal organisms (Nunez et al 2021), balancing selection would tend to retain additional diversity throughout the genomic region. It is possible that this locus reflects a complex history of introgression.

*Geukensia granosissima* (about 4 million years divergent from *G. demissa*) used to be on the Atlantic coast (Sarver et al 1992; Wares 2024) and may have localized introductions as well (Smith, White et al 2026).

That being said, the spatial pattern of amylase transitions in *G. demissa* is consistent with changes in coastal sea surface temperatures. The most notable shift in observed genotype frequencies occurs when crossing the environmental gradient associated with Cape Hatteras, NC where there is up to 5°C difference in SST over a very short coastal distance.

White (2025) showed a strong correlation between amylase genotype frequencies and latitude along the Atlantic coast (Figure 1). We will continue exploration of the amylase marker as a “genes to ecosystems” tool (Wymore et al. 2011) for salt marsh ecology.

## Acknowledgements

First and foremost, this project could not have been completed without the field and FedEx assistance from Tanya O’Reilly, Katie Miller, Davis S. Johnson, Roland Hagan, Thomas Grothues, Jenny Shinn, April Blakeslee, John Carroll, Jim Morley, Juliet Wong, Lauren Baena, Ken Halanych, and the folks at the Marine Biological Laboratory. Anderson Smith and Chloe Graham assisted in data collection. Funding for this project was from the UGA Office of the Vice President for Research and the Elaine Lutz Fund for Aquatic Biodiversity, as well as the National Science Foundation (LTER-2425396).

## Appendix

**Table A1.** World Ocean Atlas data (Reagan et al. 2024) for sea surface temperature at 1/4° resolution proximal to each sampled location.

| Site | Mean annual SST | Mean winter quarter SST | Mean summer quarter SST |
| --- | --- | --- | --- |
| Antigonish, Nova Scotia | 8.456 | 0.557 | 15.832 |
| Plum Island, MA | 10.185 | 3.313 | 15.832 |
| Woods Hole, MA | 10.359 | 4.919 | 16.63 |
| Goldstar Battalion Beach, NY | 12.415 | 4.928 | 20.106 |
| Green Island, NY | 13.138 | 4.086 | 20.991 |
| Boat Basin, NJ* | 14.69 | 4.621 | 20.921 |
| Bivalve, NJ* | 13.879 | 4.701 | 24.193 |
| George Island Landing, MD | 14.376 | 5.699 | 22.834 |
| Gloucester Point, VA | 13.594 | 7.327 | 24.719 |
| Roanoke Sound, NC | 16.731 | 7.493 | 26.435 |
| Atlantic Beach, NC | 21.722 | 12.812 | 26.878 |
| Wilmington, NC | 19.31 | 11.711 | 27.122 |
| Charleston, SC | 18.397 | 11.879 | 26.366 |
| Savannah, GA | 21.646 | 12.263 | 28.19 |

